# Tracking electron uptake from a cathode into *Shewanella* cells: implications for generating maintenance energy from solid substrates

**DOI:** 10.1101/116475

**Authors:** Annette R. Rowe, Pournami Rajeev, Abhiney Jain, Sahand Pirbadian, Akihiro Okamotao, Jeffrey A. Gralnick, Mohamed Y. El-Naggar, Kenneth H. Nealson

**Affiliations:** Department of Earth Sciences, University of Southern California, Los Angeles, CA, 90089; Department of Microbiology and Biotechnology Institute, University of Minnesota, St. Paul, MN, 55108; Department of Physics and Astronomy, University of Southern California, Los Angeles, CA, 90089; Global Research Center for Environmental and Energy based on Nanomaterials Science, National Institute for Materials Science (NIMS), Tsukuba-city Ibaraki, Japan, 305-0047; Department of Biological Sciences, University of Southern California, Los Angeles, CA, 90089; Department of Chemistry, University of Southern California, Los Angeles, CA, 90089

**Keywords:** *Shewanella*, Extracellular electron transfer, Cathode oxidation, Reverse electron flow, Electrosynthesis

## Abstract

While typically investigated as a microorganism capable of extracellular electron transfer to minerals or anodes, *Shewanella oneidensis* MR-1 can also facilitate electron flow from a cathode to terminal electron acceptors such as fumarate or oxygen, thereby providing a model systems for a process that has significant environmental and technological implications. This work demonstrates that cathodic electrons enter the electron transport chain of S. *oneidensis* when oxygen is used as the terminal electron acceptor. The effect of electron transport chain inhibitors suggested that a proton gradient is generated during cathode-oxidation, consistent with the higher cellular ATP levels measured in cathode-respiring cells relative to controls. Cathode oxidation also correlated with an increase in the cellular redox (NADH/FMNH_2_) pool using a bioluminescent assay. Using a proton uncoupler, generation of NADH/FMNH_2_ under cathodic conditions was linked to reverse electron flow mediated by the proton pumping NADH oxidase Complex I. A decrease in cathodic electron uptake was observed in various mutant strains including those lacking the extracellular electron transfer components necessary for anodic current generation. While no cell growth was observed under these conditions, here we show that cathode oxidation is linked to cellular energy conservation, resulting in a quantifiable reduction in cellular decay rate. This work highlights a potential mechanism for cell survival and/or persistence in environments where growth and division are severely limited.

## Introduction

Microbes possess an impressive diversity in the types of oxidation and reduction reactions they perform to conserve energy. While generally thought of in the context of growth, in low energy environments (where electron donors and/or acceptors become limiting) respiration may be solely utilized for maintenance (1, 2)—a state difficult to study with traditional culturing methods and approaches to studying microbial physiology. The energy acquired, and/or the proportion of that energy used for growth vs. maintenance, are difficult to quantify in natural systems, especially when solid substrates are utilized in energy acquisition. Newly developed electrochemical approaches have been used to better understand mineral respiring microbes, resulting in quantitative measurements of electron flow in microbes capable of utilizing solid substrates and/or electrochemically active mediators (3). For example, mineral-reducing organisms will reduce anodes in place of insoluble terminal electron acceptors such as manganese or iron oxides (3).

The mechanisms of extracellular electron transfer (EET) from the cellular interior to external electron acceptors are best understood in the *Gammaproteobacteria Shewanella oneidensis* strain MR-1. Under anaerobic conditions, with an organic acid electron donor and in the presence of a suitable sink for electrons on the cell exterior, electrons from the MR-1 inner membrane quinone pool are transferred to the inner membrane linked tetraheme cytochrome CymA (4, 5). Electron transfer to the cell exterior is thought to depend on protein-protein interactions between CymA with periplasmic electron carrying proteins such as the small tetraheme cytochrome (Cct) or the flavocytochrome fumarate reductase FccA (6-8). Cct and FccA likely interact with the Mtr EET respiratory pathway through MtrA, a periplasmic decaheme cytochrome (8). MtrA helps traffic electrons across the outer membrane via interactions with the MtrB porin and with decaheme lipoprotein cytochromes localized to the exterior of the outer membrane (MtrC, OmcA etc.) (9). These complexes (illustrated in the *Supporting information* Fig. S1A) have been shown to be involved in electron transfer (either directly or indirectly) to solid substrates such as solid state electrodes, and manganese or iron (oxy)hydroxides (10).

In addition to anode reduction, it has been demonstrated that mineral-reducing microbes like *Shewanella,* can also facilitate cathodic reactions—transferring electrons from an electrode to a more oxidized terminal electron acceptor (11-15). Under anaerobic conditions in MR-1, this process can be coupled to fumarate reduction and has been proposed to result from a reversal of the electron transport pathways that function in anode reductions (16, 17). However, the potential for energy conservation remains unclear, especially given the relatively small energetic gains from coupling the Mtr pathway to anaerobic terminal electron acceptors (16). Coupling cathode oxidation with oxygen reduction has been observed previously in other organisms (12, 14), though never specifically reported in MR-1. The oxygen couple thermodynamically allows a higher energy gain, though it is unknown whether MR-1 cells are able to capitalize on electrons from an extracellular source to generate a proton motive force (PMF). Given the highly enriched cytochrome network in *Shewanella,* it is plausible that non-specific reduction reactions could be occurring between cytochromes and oxygen resulting in a catalytic reduction of oxygen without PMF generation. Alternatively, reversing the EET pathway may result in electrons entering the cellular quinone pool and/or interacting with one or more of the inner membrane cytochromes in a way that allow electrons to flow to one of the three terminal oxygen reducing cytochrome reductases (i.e. *cbb*_3_, *aa*_3_, or *bd*) (18).

To better understand energy conservation by MR-1 under cathodic conditions, we used an electrode to impose electron donating redox potentials in an aerobic environment lacking exogenous organic carbon sources. Under these conditions, we set out to understand: 1) whether or not electrons from a cathode that enter MR-1 can be utilized for cellular energy conservation and, 2) the pathways involved in electron flow from a cathode to oxygen. Understanding the physiology behind these biologically-mediated cathodic processes, may allow us to optimize and/or utilize microbes for various microbe-electrode applications such as electrosynthesis, as well as to better understand microbial physiology under a variety of redox conditions.

## Results

### Electrons flow from a cathode to the S. *oneidensis* MR-1 cellular electron transport chain

Oxygen reducing cathode conditions were investigated in three-electrode electrochemical cells using indium tin-doped oxide (ITO) coated glass working electrodes poised at -203 to -303 mV vs. SHE and covered with a monolayer biofilm. Significantly more cathodic current was generated compared to control conditions (abiotic/cell-free media, and killed cell biomass), and this effect was linked to the presence of oxygen in MR-1 monolayer cathode biofilms *(Supporting information* Fig. S2). Comparing the rates of electron flow between anodic (+397 mV, anaerobic 10 mM lactate) and cathodic conditions (-303 mV, aerobic, no exogenous electron donor) for the same monolayer biofilms, cathodic conditions yielded 23.7 ± 5 times more current production on average (n = 4). The possibility of these cathode derived electrons entering the cellular electron transport chain (ETC) was investigated in this system using the combination of a redox active dye (RedoxSensor Green™ [RSG]) and ETC inhibitors. RSG is a lipid soluble redox active dye, previously shown to fluoresce in actively respiring aerobic and anaerobic microbial cells (19–21).

RSG fluoresces green when reduced and active accumulation of the reduced dye can be linked to respiratory conditions (i.e. active electron flow through the ETC). While the specific oxidoreductases involved in RSG reduction are not known, previous work in MR-1 investigated the effects of ETC inhibitors during aerobic growth on lactate and oxygen (19). Inhibition of RSG activity was seen when an ETC inhibitor was utilized; specifically it was noted that inhibition of electron flow downstream of the terminal oxidase step also interfered with the cellular reduction of RSG (19). In this work we were able to quantify an increase in RSG fluorescence in MR-1 cells under applied cathodic potential (-303 mV vs. SHE) in the presence of oxygen. Minimal fluorescence was observed under open circuit conditions. The fluorescence signal seen under cathodic conditions was strongly inhibited by potassium cyanide, a respiration inhibitor (Fig. 1). These observations support the notion that active cellular electron flow is required for active RSG reduction and accumulation in cells. Time lapse videos from these experiments demonstrate the rapid nature of this process; a marked increase in RSG signal intensity can be observed within 15 minutes (three frames) of applying a cathodic potential (Videos S1-S3). Additionally, RSG fluorescence significantly decreased within 15 min after addition of potassium cyanide, an inhibitor of cytochrome C oxidase (Fig. 1, Videos S1-3), similar to the time frame under which RSG signal dissipates in formalin-killed cells according to the manufacturer (Molecular Probes, Life Technologies). Cathodic current was also mitigated by cyanide addition (Fig. 2), supporting a requirement for terminal cytochrome C oxidases to facilitate electron flow from a cathode. Removal of cyanide from a cathode biofilm after 30 min exposure allowed for recovery of both cathodic current (68% of wild-type current recovered) and RSG fluorescence suggesting this effect is due to the reversible inhibition of cytochrome C oxidases.

**Fig. 1.**
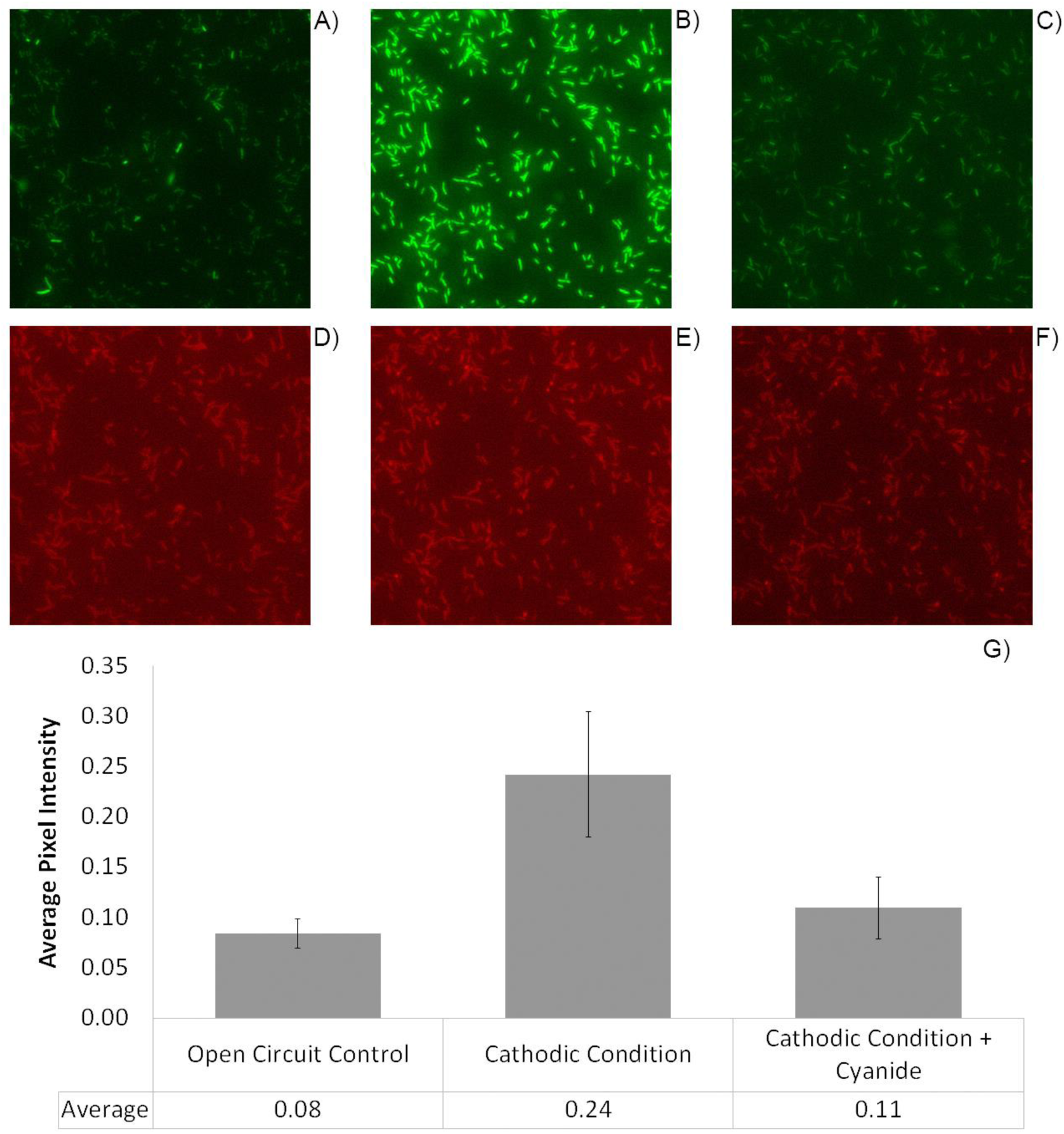
Representative images of MR-1 cells attached to ITO-coated glass and treated with RedoxSensor™ Green (A-C) and the lipid stain FM 4-64FX (D-F) as described in materials and methods. Fluorescence intensity is compared between the control (open circuit A & D), cathodic conditions (-303mV vs. SHE B & E) and cathodic conditions with an inhibitor of cytochrome C oxidase added (-303mV vs. SHE with 5mM KCN added C & F). Average pixel intensity per cell was calculated for approximately 80 images for six time points per condition (average cells per image n =2271, 1904, 2234 for control, cathodic, and inhibition conditions respectively). Error bars indicated standard deviation in average pixel intensity (per population) per image (80 images analyzed per experimental condition).

**Fig. 2.**
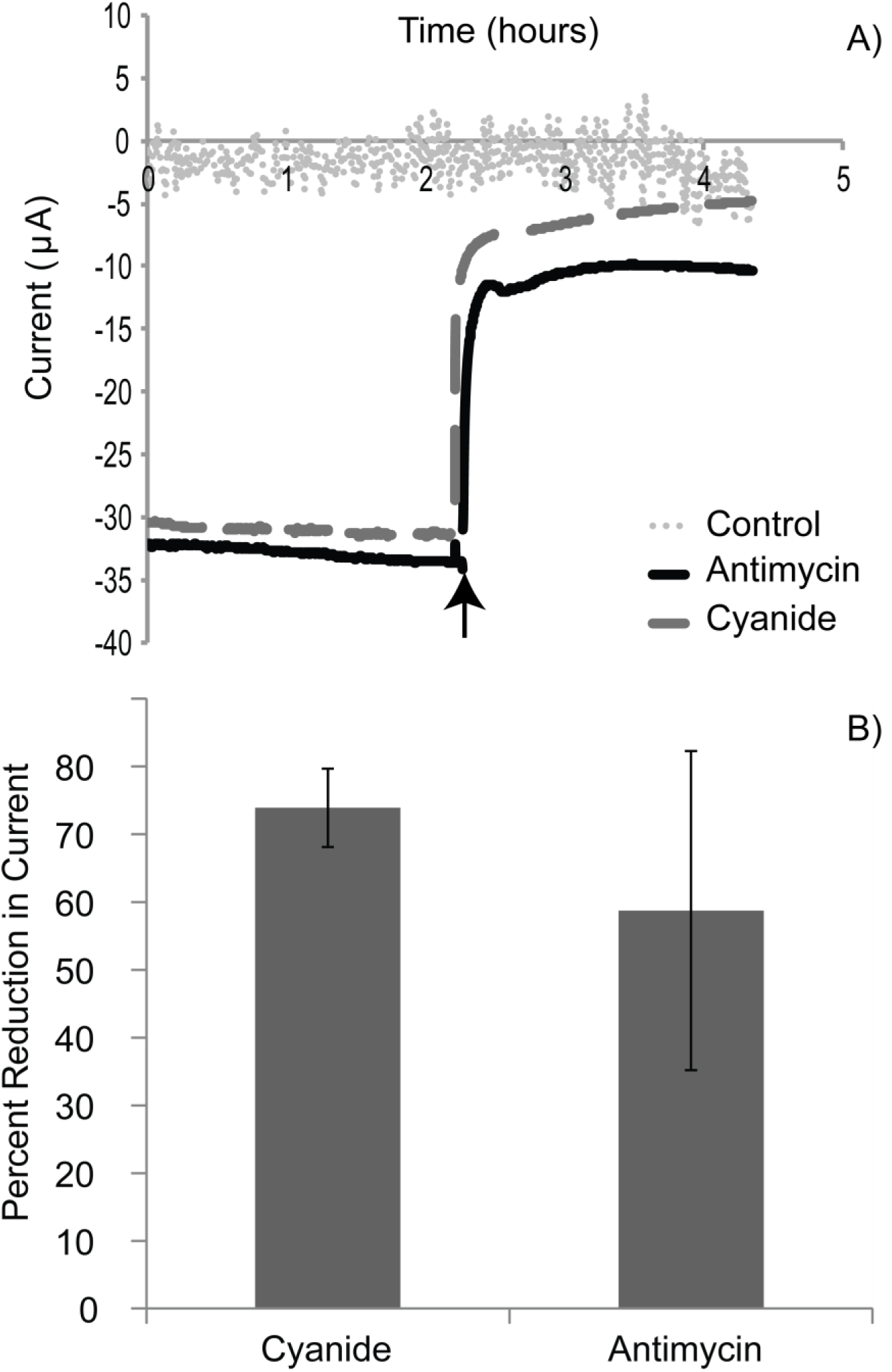
Example chronoamperometry plots for MR-1 cells attached to a cathode (one shown of n = 3) and a cell-free control electrode (-303mV vs. SHE) with the electron transport chain inhibitors potassium cyanide (5 mM) and Antimycin A (20 μM) added (addition indicated by arrow), which inhibit cytochrome C oxidase and quinone oxidoreductases respectively (A). Control sample treated with both potassium cyanide and Antimycin A at time indicated. The average percent reduction in cathodic current (average of negative current production over hour pre and post injection) when electron transport chain inhibitors added is illustrated for n = 3 reactors (B). Error bars represent one standard deviation of triplicate experiments.

While cyanide is a general cytochrome oxidase inhibitor, and several cytochrome oxidases have been shown to pump protons (22), inhibition at quinone proton translocation sites was also tested. Addition of Antimycin A (an inhibitor of quinone oxidoreductases) also resulted in rapid and marked decrease in cathodic current while having no effect on abiotic controls (Fig. 2). Notably, a very large inhibition of current (60-70%) was observed within the first minute of inhibitor addition (Fig. 2). RSG activity was monitored in cathode biofilms under all inhibitor experiments. While a dissipation of fluorescence can still be observed using Antimycin A (Videos S4-S6), the quantification of RSG post addition was made difficult by autofluorescence of Antimycin A. Though Antimycin A, as a quinone mimic may interact non-specifically with other quinone oxidoreductases, it has been shown to preferentially inhibit oxidation of ubiquinone by the cytochrome bc1 complex in mitochondria (23). These results demonstrate that electron flow from a cathode passes through at least one coupling site in the cellular electron transport chain.

### Cellular energy carrier quantification in cathode oxidizing S. *oneidensis* MR-1 cells

To further investigate if a proton gradient is generated under cathodic conditions and can consequently result in generation of cellular energy carriers, we measured pools of ATP and ADP within cathode biofilms. We compared the difference in ATP to ATP+ADP ratios (to normalize for differing overall cellular nucleotide levels) for replicate biofilms exposed to either cathodic conditions (-303 mV vs. SHE), cathodic conditions treated with the protonophore uncoupler carbonyl cyanide m-chlorophenyl hydrazine (CCCP), and poised potential conditions (197 mV vs. SHE) where minimal anodic current flow was observed (average of five replicates 0.19 ± 0.4 μA). This minimal anodic current was a means of accounting for the background heterotrophy and/or cellular energy obtained from storage products in this carbon free system, although it was impossible to control for the potential for additional electron equivalents added in this system—specifically from decaying cell biomass, or hydrogen produced on the counter electrode. Nonetheless, a significantly higher ATP/ADP+ATP ratio was quantified under cathodic conditions compared with either control condition, though significant variation was observed under the minimal anodic regime (Fig. 3). While we did not quantify AMP levels, the ATP to ATP + ADP ratios also supported an increase in energy charge state of the cells under cathodic conditions.

**Fig. 3.**
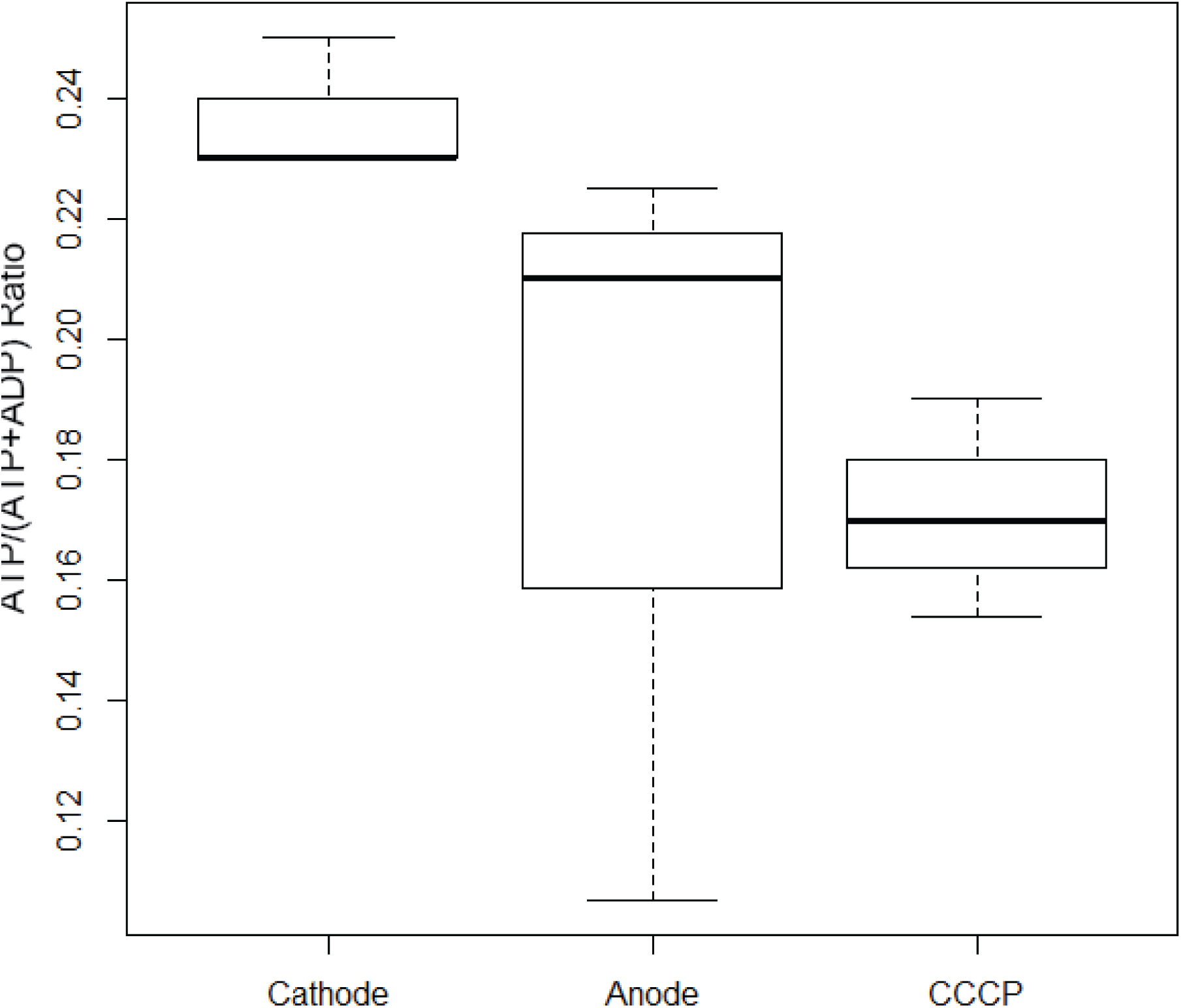
Comparison of ATP levels normalized to combined ATP + ADP recovered in MR-1 biofilms maintained under poised electrode conditions (-303 or 197 mV vs. SHE) for 24 hours. The conditions compared reflect cathodic current generation with an electrode potential poised at -303 mV (Cathode [n=9]), minimal anodic current generation with an electrode potential of 197 mV (Anode [n=9]), and cathodic current generation as described above with a three hour CCCP treatment prior to ATP recovery (CCCP [n=9]). All treatments performed under exogenous-carbon free and aerobic environmental conditions. Box plot reflects the median value (black bold line), one standard deviation around the mean (white box), and the data spread (dashed line).

Statistically significant cell loss was observed in open circuit controls *(Supporting information* Fig S8), likely due to a lack of applied potential or energy input required to maintain cell biomass on the electrode. The minimal anodic condition serves as a similar environmental condition to the cathode treatment (same media and oxygen inputs) with the exception the direction of electron flow on the electrode differs and, unlike open circuit controls, cell biomass does not statistically change throughout the course of the experiments (2.2x10^7^ ± 5×10^6^ and 2.3×10^7^ ± 1.2×10^7^ cells per biofilm for cathode and minimal anode respectively). The per cell ATP values estimated in this work fall between 0.13 and 0.68 fmol of ATP per cell. Though cellular ATP levels can vary across microbes as well as growth rates (shown to range six orders of magnitude across taxa, (24)), the per cell ATP levels observed in this work were similar to environmentally sampled *Escherichia coli* cells (0.18 - 0.25 fmol per cell) (24, 25).

The observed difference in ATP/ATP+ADP ratios between MR-1 wild type cathodic biofilms and 3-hr CCCP-treated MR-1 cathodic biofilms (Fig. 3), supports the notion that MR-1 generates a proton gradient during cathode oxidation. Depending on the degree of proton accumulation, this PMF could also aid in reverse electron flow—electron flow from the generally higher potential quinone pool to oxidized cellular electron carriers—resulting in the generation of cellular reducing equivalents (i.e. NADH) from cathodic electron flow (model illustrated in *Supporting information* Fig. S1B). To directly test whether cathodic electron flow can be converted to cellular reducing power in the form of NAD(P)H and/or FMNH_2_, we used the bacterial luciferase enzyme, inserted downstream of the *glmS* gene into a neutral site in the bacterial genome via transposition with the *lux* operon *(luxCDABE)* (26), as a real time *in vivo* marker for the cellular redox pool.

The Lux enzyme system performs a well-characterized cytoplasmic process that generates light using an oxygen molecule, reduced flavin mononucleotide (FMNH_2_) and an activated (via NADPH & ATP) aldehyde functional group (27). Though, there are multiple factors that influence light production, the reducing pool has been directly linked to light production (28-30). Expression of the *lux* operon is constitutively driven by the p1 promoter, resulting in light production under aerobic growth conditions. The variation of enzyme levels, depending on the growth history of the cells, makes it difficult to compare light production across different experiments where the energy investment in protein production vary. However, by comparing across cell populations with similar growth histories and maintaining non-limiting oxygen and aldehyde concentrations for the Lux reaction we are able to link lactate concentration (which in turn affects electron flow and the redox pool) to light production in normalized cell populations *(Supporting information* Fig. S3).

Light production was limited for aldehyde under cathodic conditions (no exogenous carbon), as demonstrated by the rapid increase in light production when 0.002% decanal was provided (modest initial increase likely due to aldehyde diffusion across the cell membrane), peaking 0.5 hours post decanal addition (Fig. 4). Increased light production does not appear to be based on decanal conversion to reducing power as no light was observed under the control (minimal anodic current condition), and MR-1 did not grow aerobically with decanal as the sole carbon source. Though it was difficult to compare light intensities across various experiments, the trend of increasing light production with increasing negative current production compared to controls was consistently observed and was independent of the order of poised potential conditions *(Supporting information* Fig. S4). We also noted that the magnitude of current generated was positively correlated with light production *(Supporting information* Fig. S4). Given the variety of potential sources of reducing equivalents in MR-1 (cellular storage products, endogenous cell decay, etc.), it is difficult to determine if there is direct link between cathodic electrons and the cellular reducing pool (total cellular electron carriers) from these data alone. However, the increase in the cellular reducing pool observed in *SO-lux* strains, along with the likelihood of PMF generation from cathodic electron flow suggest that reverse electron flow could be operating in MR-1 cells under cathodic conditions.

**Fig. 4.**
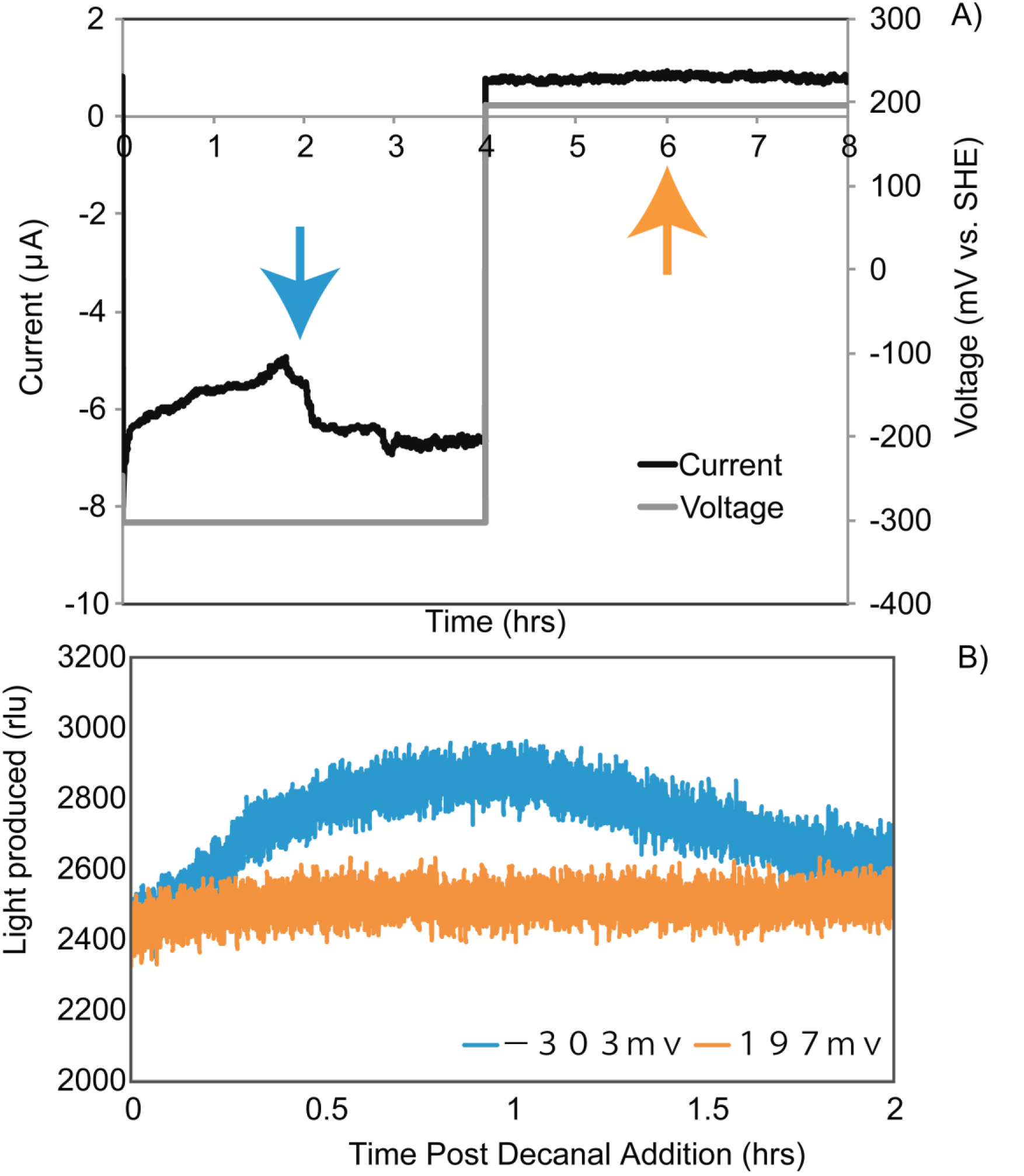
Representative demonstration (one of four) of *SO-lux* electrode-biofilm current (A) and light (B) production for an electrode poised at a cathodic current generating redox potential (−303 mV SHE, 0 to 4 hours) and then switched to a non-cathodic redox potential (197 mV SHE, 4 to 8 hours) where minimal anodic current is observed. Decanal is added at a concentration of 0.002% at two hours into each poised potential incubation (as indicated by the colored arrows). Light production quantified by a photon multiplying tube presented in relative light units (RLU) is depicted for the two-hour period post decanal addition (B)—corresponds to the 2-hour period following blue and orange arrow in panel A.

### Reverse electron flow enhances the cellular reducing pool in S. *oneidensis* MR-1

Reverse electron flow involves utilizing a proton gradient to drive electrons from a higher potential state (i.e. quinone pool) to a lower potential state (i.e. NAD+) in the cellular electron transport chain (31, 32). Under conditions of reverse electron flow, generation of NADH can be inhibited by a protonophore uncoupler (33). To determine if cathodic electron flow resulted in an enhanced redox pool through reverse electron flow, the protonophore uncoupler CCCP was used in combination with *SO-lux* strain to determine if dissolution of the inner membrane proton gradient affected light generation via the luciferase enzyme—utilizing light as an intracellular marker for the cellular redox pool. Addition of CCCP to *SO-lux* showed a marked (nearly order of magnitude) decrease in light production (Fig. 5A), though no statistically significant effect was observed on current *(Supporting information* Fig. S5). The inhibitory effects of CCCP on light production were mitigated by addition of lactate under otherwise identical cathodic conditions *(Supporting information* Fig. S6). This demonstrates that the decline in light production resulting from CCCP addition requires a limitation in NADH, or a more oxidized reducing pool to allow for reverse electron flow. It also argues against CCCP directly affecting NADH oxidation rates, as the same collapse in light production would be expected in the presence of lactate.

**Fig. 5.**
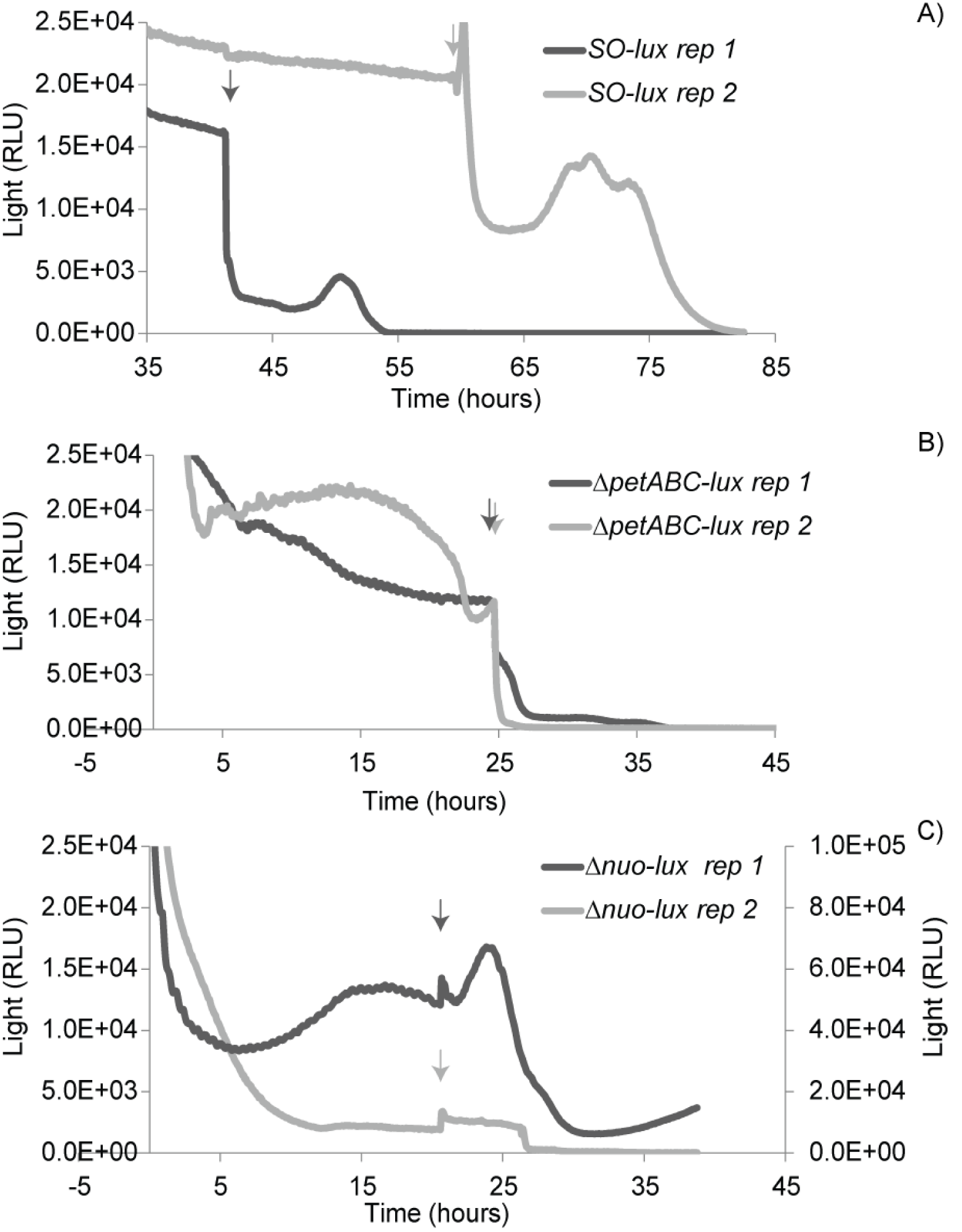
Light quantification in cathode biofilms poised at -303 mV vs. SHE for two (of n = 3) replicates of MR-1 strains amended with the *lux* operon: wild-type MR-1 (*SO-lux* [A]), the *bc*_1_ complex mutant (Δ*petABC-lux* [B]), and the Complex I mutant (Δ*nuo-lux* [C]). After at least 20 hours under cathodic conditions and at the time points indicated by arrows on each plot, the protonophore uncoupler CCCP was added to each reactor. Light was measured via photon multiplier tube and presented in relative light units (RLU). Plots depict 45 to 50-hr period around CCCP addition. Replicate 1 for Δ*nuo-lux* demonstrated 2-fold higher light intensities and is therefore plotted on a larger light intensity scale (right axis in plot C).

A similar decline in light production was observed when CCCP was added to a mutant of S. *oneidensis* lacking the *bc*_1_ complex of the electron transport chain amended with the *lux* operon (Δ*petABC-lux*) (Fig. 5B).The *bc*_1_ complex is not essential for aerobic respiration (34) as MR-1 has multiple ETC routes to reduce oxygen. However, the PetABC complex has the potential to maximize PMF generation under aerobic conditions (i.e. Q-cycle vs. Q-loop) (33). Additionally, the observed loss of light production upon CCCP addition suggests that PetABC is not involved in mediating electron flow to the cellular redox pool in MR-1 as has been suggested in other organisms (35, 36). Unlike other mutants, the Δ*nuo-lux* strain (Complex I deletion in S. *oneidensis* amended with *lux),* did not demonstrate the same initial decline in light production when treated with CCCP (Fig. 5C). The observation that in the absence of Complex I PMF does not have the same effect on cellular light production/NADH levels, implicated the MR-1 Complex I in reverse electron flow. This result could be recreated in the wild type *SO-lux* strain when the Complex I specific inhibitor Piericidin A was added to a cathodic biofilm. Piericidin A addition prevented the decrease in light production observed in the wild-type cells alone, further supporting a Nuo role for maintaining the redox pool under cathodic conditions in MR-1 *(Supporting information* Fig. S6).

CCCP also did not affect light production under the control condition where a minimal oxidizing potential was poised and a minimal anodic current was observed (average 23 ± 1 and 34 ± 3 nA) *(Supporting information* Fig. S6). This supports the importance of cathodic electron flow in addition to an oxidized redox pool and a reversible proton-translocating NADH/Ubiquinone oxidoreducase for driving the formation of redox pool linked to PMF and the subsequent collapse as shown by the Lux reaction under these electron donor-limited conditions.

### Cathode oxidation results from a reversal of multiple extracellular electron transport routes in S. *oneidensis* MR-1

In order to assess the route of electron flow into cells under cathodic conditions, we utilized a set of gene deletion mutants of various extracellular electron transport pathways, as well as inner membrane electron transport chain mutants (Table 1, illustrated in the *Supporting information* Fig. S1). Of the mutants tested, the mutant deficient in all three of the terminal cytochrome oxidases (Δcox-all) demonstrated the greatest percentage decrease in cathodic current production, though the individual cytochrome mutants displayed near wild-type cathodic current levels (Table 1, *Supporting information* Fig. S7). Deletion of all five outer membrane multiheme cytochromes (Δ*omc*-all mutant) also significantly decreased cathodic current production—only slightly greater than the Δ*mtrC/omcA* double mutant (Table 1, *Supporting information* Fig. S7). Mitigated cathodic current production was observed in the majority of the other cytochrome mutants tested, including a variety of periplasmic electron carriers. Surprisingly, this included proteins like DmsE; a periplasmic decaheme cytochrome shown to be associated with DMSO reduction (37). Some cathodic current generation could be recovered when a copy of the *dmsE* gene behind the ribosomal binding site of *mtrA* was added to the Δ*dmsE* mutant in trans, suggesting the DMSO respiratory pathway is important in cathodic electron flow. Coupled with the observations that deletion of *dmsB* and/or the entire *dms* operon (Δ*dms*-all) significantly decreases cathodic current generation (and to similar levels of the Δ*omc*-all mutant), there is the potential that multiple extracellular electron transport pathways are running in parallel and/or cooperatively during cathode oxidation. It is difficult, however, to distinguish the hypothesis of parallel pathways of cathodic electron flow from the possibility of differential gene expression affecting cathodic current generation among the various mutants.

Cyclic voltammetry (CV) was performed on all mutant strains utilized in these experiments. The only two mutants to demonstrate similar CV profiles for aerobic cathodic electron flow to wild type (Fig. 6) were the Δ*fccA* and Δ*ccoO* (data not shown). The other mutants tested showed a significantly decreased onset potential and magnitude of cathodic electron flow that was only slightly enhanced from controls (data not shown); consistent with the observed decrease in current generated during chronoamperometry experiments for these mutants. Interestingly, a 180-200 mV increase in onset potential was observed when comparing MR-1 aerobic/oxygen-reducing cathodic biofilms and MR-1 anaerobic/fumarate-reducing cathodic biofilms (Fig. 6). This result is still consistent with a model of outer membrane multiheme cytochromes like MtrC and OmcA being involved in cathodic electron flow, if one assumes a model of flavin-binding under anaerobic conditions and not under aerobic conditions as has been previously reported (17). Under aerobic conditions we observed a 0 to 20 mV vs. SHE cathodic current onset potential, whereas under anaerobic conditions with fumarate, an onset potential of -160 to -180 mV vs. SHE was observed (Fig. 6) which match the corresponding potentials proposed for the outer membrane cytochromes with unbound and bound flavins respectively (38). These observed redox potentials further support the role of outer membrane cytochromes in cathode oxidation, though it does not exclude the possible activity of other pathways with similar oxidation/reduction potentials.

**Fig. 6.**
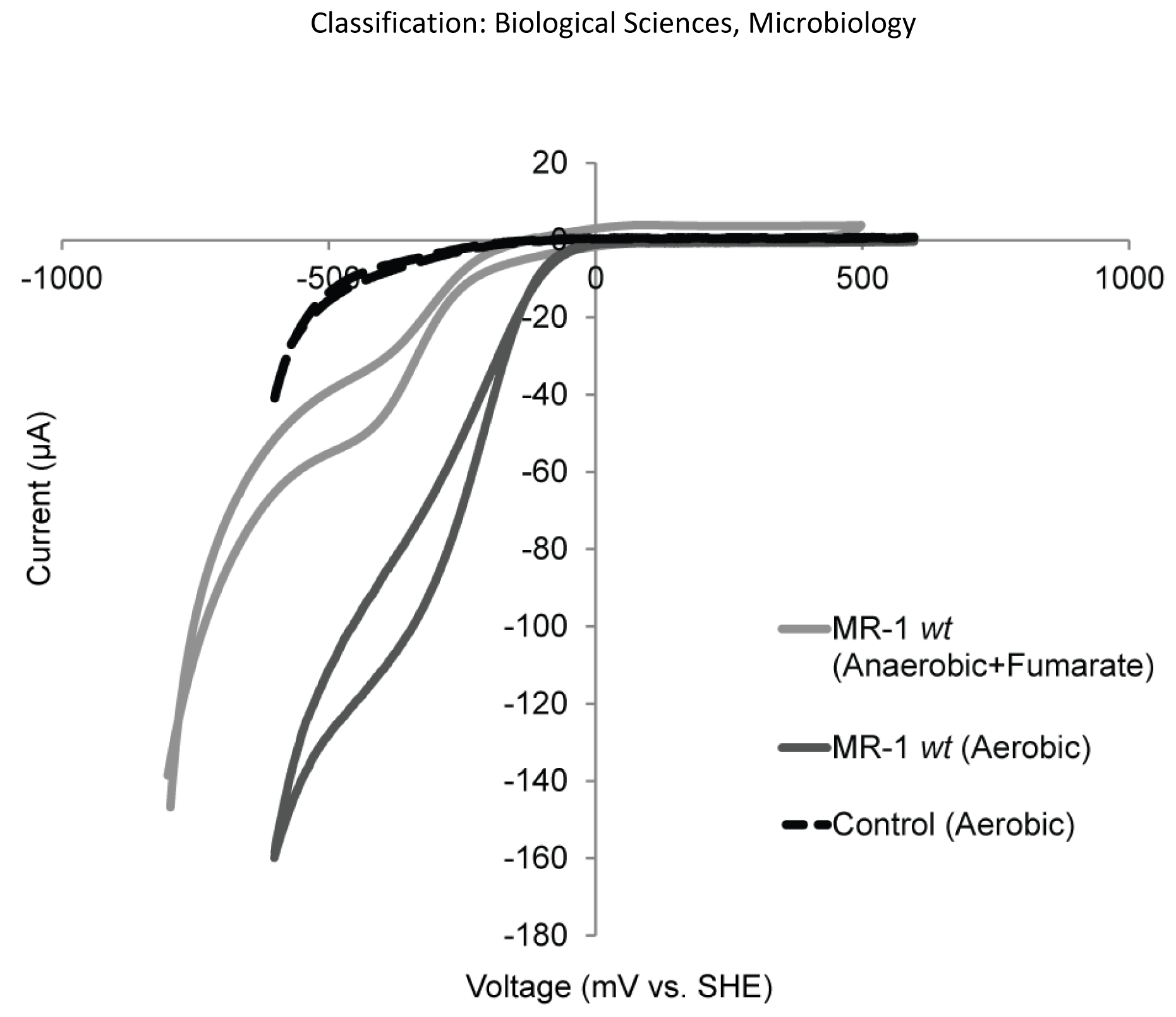
CV curves taken for MR-1 cathodic biofilm poised at -303 mV vs. SHE incubated under aerobic conditions with oxygen as a terminal electron acceptor or anaerobic conditions with fumarate as a terminal electron acceptor (17). CV (one of n = 3 shown) for cell-free abiotic control under aerobic conditions (dashed line) also shown. Scans were run at 5 mV/s for aerobic electrodes and 10 mV/s under anaerobic conditions.

### Implications for cathode-oxidation on minimizing cellular decay

As expected, cell growth was not observed over a five-day period under cathodic conditions (*Supporting information* Fig. S8A). However, cell decay (loss of cell biomass through death and lysis) was also not observed under cathodic conditions. Open circuit conditions demonstrated a statistically significant loss in surface attached cell biomass over the same experimental period (approximately 17% cell loss per day, *Supporting information* Fig. S8). This survival stands in sharp contrast to the cellular decay rate of planktonic cells under comparable conditions (identical medium and similar starting OD), where a 48% loss of biomass is expected over the same time period *(Supporting information* Fig. S8). While it is difficult to distinguish cell detachment from cell decay under biofilm conditions, over a five-day period the loss of cell biomass observed in the minimal anodic current condition is consistent with the amount of cell loss observed under open circuit conditions (~ 4-day half-life) though not statistically significant within a 24-hour period *Supporting information* Fig. S8). These observations add further support to the capacity for cellular energy conservation under cathode oxidizing conditions, and imply that this energy could be harnessed to provide cell maintenance to stave off cell death and decay and/or maintain cellular attachment in a biofilm.

## Discussion

This report links cellular energy conservation with cathode oxidation coupled to oxygen reduction in S. *oneidensis* MR-1. The periplasm and outer-membrane of MR-1 is enriched in redox active cytochromes, raising the possibility that electrons from an exterior cathode could be routed along this redox network to a terminal oxidant in a biocatalytic process, rather than interact with inner-membrane components that would allow cells to conserve energy. In the case of an oxygen-reducing MR-1 cathode biofilm, this work demonstrates that electrons interact with coupling sites of the cellular inner membrane electron transport chain suggesting the generation of PMF is feasible. As a result, quantifiable differences in the cellular ATP pools and NADH levels between cathode-respiring electrode-attached cells and various controls.

The differences in proportion of reduced cytoplasmic electron carriers in MR-1 under cathodic conditions were linked to reverse electron flow by observing the effect of protonophore uncoupler under NADH-limiting conditions (i.e. electron donor limited) and in the presence of Complex I. Specifically, addition of CCCP caused a decrease in light production, most likely reflecting the loss of NADH generation from reverse electron flow which requires PMF. An alternative hypothesis for this observation is an increased rate of NADH oxidation due to removal of the PMF as might be expected under certain aerobic and heterotrophic conditions where NADH was being oxidized by the cellular ETC. However, the collapse in light production was prevented under cathodic conditions when an electron donor (lactate) was provided and/or when the activity of Complex I was impaired. In both of these cases, reverse electron flow would be inhibited as NADH generation is dependent both on a high NAD+/NADH ratio and the enzymatic ability to utilize PMF for NADH generation. If NADH oxidation by the ETC alone was driving this effect, additional NADH should not mitigate light production, nor should the absence of the Complex I as MR-1 has alternate NADH/quinone-oxidoreductases (i.e. *ndh*) (39, 40). It is worth noting that under cathodic condition, electron flow through the ETC is likely replete given that the magnitude of the cathodic current is 20 fold higher than the current produced by the same biofilm under anodic conditions. If NADH oxidation is competing with the cathode for the ETC, as would be expected if PMF collapse increased NADH oxidation, cathodic current would also likely be inhibited. However, upon CCCP addition, cathodic current either does not change or marginally increases.

Reverse electron flow, as has been described in chemolithoautotrophs, results in the generation of NADH (41). Implicate for cellular biosynthesis and/or carbon fixation is the conversion of NADH to a variety of cellular electron carriers utilized in anabolism—most commonly NADPH (31). Light production via luciferase has been shown to be directly dependent on FMNH_2_ levels and indirectly on NAD(P)H levels (27), as NAD(P)H is required for both the activation of the aldehyde moiety (LuxCE). Additionally, cellular reducing equivalents (i.e. NADH, NADPH) are involved in the regeneration of FMNH_2_ via endogenous cellular enzymes (28, 42–44). While the luciferase reaction is not a quantitative marker of NADH levels specifically, the effect of CCCP on the cellular reducing pool can be observed via luciferase supporting to connectivity and or conversion of cellular reducing equivalents for biosynthesis, a process further supported by the observation of cell maintenance under cathodic conditions.

Though we could demonstrate the generation of cellular energy carriers in this system, there was no evidence of cell growth/division under these conditions. No growth was expected given that these experiments were performed in a system that lacked exogenous carbon inputs and several growth factors (i.e. vitamins and amino acids) that could provide carbon (45), and the genome contains no known complete carbon fixation pathway (46). However, the energy conserved in the form of ATP and cellular reducing power could plausibly support cell maintenance under cathodic conditions. As cathode biomass was maintained past the expected cell decay rates, it seems likely that energy acquired is being invested towards cellular upkeep. This is further supported by previous observations in MR-1 that demonstrated *Shewanella* requires energy in order to maintain cell surface attachment (47), and as such energy acquired under cathodic conditions could also be devoted to maintaining attachment. These observations support the proposal that MR-1 cells conserve energy under the studied cathodic conditions; energy that prolongs cell survival and/or allows for maintaining cell attachment.

Anaerobically, a reversal of the Mtr pathway was shown to be important for fumarate reduction from a cathode using mutants (16), but energy conservation under these conditions was not tested. However, it is unknown whether cathodic electron flow makes it to the menaquinone pool as a result of CymA interactions, though this was previously suggested through use of a menaquinone biosynthesis mutant (16). FccA has been further implicated in the transfer of electrons to the periplasmic decaheme cytochromes such as MtrA under certain conditions (8), providing a more direct route from MtrA to FccA under cathodic conditions. The relatively low redox potential window available under these anaerobic conditions also makes it unlikely that MR-1 is able to capitalize on these redox couples to drive proton pumping—redox potentials lower than known coupling sites and overall energy yield relatively low. Conversely the energetics for aerobic cathode conditions are decidedly better owing to the greater reduction potential of oxygen as the terminal electron acceptor and the ability of cells to capitalize on the electron transport chain commonly used for aerobic respiration.

Though a model for electron flow has been developed for MR-1 anaerobic cathode oxidation using fumarate as an electron acceptor (16), much less was known about electron flow under cathodic conditions using oxygen as a terminal electron acceptor. Our results highlight that the network of proteins might be more complicated and the detailed route(s) at this time remain speculative. Although the use of gene deletion mutants has allowed us to highlight several possible pathways, the degree of decrease in cathode-oxidizing activity in non-redundant pathways suggest that either: 1) the architecture of the cytochrome network is in some way cooperative and/or plays an important functional role in the rate of cathodic electron flow; or 2) differential gene expression across mutants is affecting cathode oxidation activity.

While a specific pathway has not been identified aerobically, overlapping proteins have been highlighted as being involved under both aerobic and anaerobic conditions—namely the Mtr pathway. However, analysis via cyclic voltammetry suggests a distinct redox difference in the route of entry from MR-1 cells under aerobic conditions. Under anaerobic conditions, a flavin-bound cytochrome intermediate was proposed to be important for cathode oxidation under anaerobic conditions, driving the observed low redox potentials (17). It has recently been proposed that flavin-binding is mediated by a disulfide bond formed under aerobic conditions in domain three of several MtrC homologs including OmcA and MtrF, which is thought to prevent flavin binding to these outer membrane cytochromes in the presence of oxygen (48). Our results are consistent with cathodic activity under aerobic conditions being mediated by a non-flavin bound c-type cytochrome (due to the presence of oxygen), as the redox potentials we observe are more in line with those reported for Mtr proteins lacking flavins (9, 49). While these non-flavin bound cytochromes likely play a role in cathode oxidation, our data suggest that the canonical MR-1 outer membrane cytochromes (MtrC and homologs) are not the only available path for electrons.

The ability of *Shewanella* to conserve energy under aerobic cathode-oxidizing conditions may have important implications for the utility of this organism for microbe-electrode applications. The specifics of how electrons enter the cell, and how this is coupled to a terminal electron acceptor, affects cellular energy levels, potential for growth, cell maintenance, as well as overall process rates and long term activity of these reactions. The environmental implications of these observations are more difficult to assess. In essence, are there environmental conditions under which *Shewanella* can capitalize on the reversibility of the extracellular electron transport systems for acquiring energy for cell maintenance? *Shewanella* is an environmentally ubiquitous organism that is often found in redox transition zones or complex sediments (50), it is likely that this organism is commonly faced with shifts from anaerobic to aerobic conditions. Though it appears to be a rate-adapted organism under carbon-replete anaerobic conditions (trading reaction rates for energetic efficiency), it likely has mechanisms for persistence in variable environments that are not presently well understood. Our results suggest the potential for MR-1 under carbon-limiting conditions in the presence of oxygen to capitalize on reduced minerals as electron sources (possibly ones generated previously in the absence of oxygen), for non-growth linked energy conservation. This could potentially highlight an important evolutionary advantage to such a reversible electron transport pathway. This work may also have implications for understanding electron uptake in organisms that can oxidize insoluble substrates (35, 51, 52), especially in the context of subsurface microbiology as oxic and mineral rich sediments are not uncommon in the deep-sea (53, 54). Subsurface ecosystems often support orders of magnitude more microbes than should be allowable based on current energetic models and using the available organic carbon content (55). It has been postulated that lithotrophic interactions and especially lithoautotrophs could be part of this puzzle; however when growth rates have been calculated they are remarkably slow (56). A non-growth linked lithotrophic reaction could be a mechanism of potentially sustaining or slowing cell death and decay for subsurface microbes.

### Materials and Methods

#### Bacterial Strains

The bacterial strains used in this study are listed in Table 1. In the *SO-lux* strain constructed for this work, the *lux* operon was inserted into the MR-1 genome via transposition at a neutral site via a mini-Tn7-luxCDABE-tp (26), carrying both the *luxCDABE* operon behind the broad host range constitutive *Gamaproteobacterial* promoter *P1* (derived from a dihydrofolate reductase gene (57)) and the gene for trimethoprim resistance. A transposon containing plasmid was transferred to MR-1 via conjugation from *E. coli* (WMB026 cells) along with WMB026 strain containing the PTNS3 plasmid encoding a transposase. Viable *lux* mutant strains were isolated on LB media with trimethoprim (200 mg/L) and screened for bioluminescence.

The in-frame gene deletion mutants utilized in this work were constructed and validated as described previously (58). In brief, primers were designed (listed in *Supporting information* Table S1) to amplify the flanking regions of the gene targeted for deletion. These amplicons were then ligated into a suicide vector. Post vector transfer into MR-1, strains were screened for double recombinants as described by (59).

#### Cathodic Culturing Conditions

Prior to electrochemical experiments, MR-1 was grown in batch at 30^o^C in standard media previously described including Luria broth (LB) and a defined media (DM-lactate) containing 10 mM lactate as the predominant carbon source (17). Conditioning of electrodes for chronoamperometry experiments were performed as described previously (17). In brief, DM-lactate grown MR-1 cells grown were added to electrochemical cells at an O.D. 600 nm of 0.25 in fresh medium with 10mM lactate. To induce cell attachment to electrodes, the working electrodes were poised at 397 mV vs. SHE while the reactors were purged with nitrogen to maintain anaerobic conditions. After a 20 to 30 hour incubation, planktonic cells were removed from the reactors along with remaining medium. The attached cells were rinsed three times in carbon-free DM (CF-DM) medium. Carbon-free DM lacked not only lactate, but yeast extract added to traditional DM media. Culture medium used in electrochemical experiments also lacked both trace metal and vitamin amendments for cathode experiments as these can interact abiotically with electrodes. Unless indicated, cathodic conditions were applied to rinsed attached electrode biofilms in CF-DM by poising a working electrode potential of -303 mV vs. SHE and while reactors were bubbled with room air at a continuous and constant rate (5-10 mL/min).

The following electron transport chain inhibitors were added to cathode experiments at the corresponding concentrations: 5 mM Potassium Cyanide, 20 μM Antimycin A, and 5 nM Piericidin A. Various antibiotics, Ampicillin (100 μg/mL), Kanamycin (100 μg/mL), and Trimethoprin (100 μg/mL) were used for cloning and selection marker purposes. The protonophore uncoupler CCCP was added at concentrations of 20-40 μM. During Lux cathode experiments the only alteration to the above protocol was the addition of decanal to a final concentration of 0.002% to the electrochemical cells at the time-points indicated. To ensure no deleterious effects of solvents on MR-1 cells occurred, growth curves were performed on the *SO-Lux* strain of MR-1 amended with the highest concentration of DMSO used (0.05%, for the addition of RSG), as well as 0.002% decanal, and compared to normal aerobic growth in DM-lactate (*Supporting information* Fig. S9).

#### Electrochemical Conditions

The electrochemical cell design used for these experiments was described previously (60) and were constructed in house. In brief, each reactor comprised a working electrode constructed from ITO-coated glass (SPD Laboratory, Inc., Hamamatsu Japan), a counter electrode of platinum wire, and reference electrode of Ag/AgCl in a 3M KCL solution (constructed in our lab). The main reactor type maintained a liquid volume of 10-12 mL to an electrode surface area of 3.75 cm^2^. Electrochemical cells were designed for use with purchased ITO plated glass coverslips (SPI, Westchester, PA) which had a 3.61 cm^2^ surface area to an 8 mL volume.

The majority of experiments were performed using controlled voltage conditions and measuring current production (i.e. chronoamperometry) using a potentiostat. The majority of chronoamperometry experiments were performed using eDAQ, quad channel potentiostats and the corresponding eChart software (eDAQ Inc., Colorado Springs, CO), however Lux chronoamerometry experiments were run on a Gamry 600 using Framework software (Gamry, Warminster, PA) or a VMP3 potentiostat (BioLogic Company, France) using parameters listed. Cyclic voltammetry was also performed using the Gamry 600 or the VMP3 potentiostats using the parameters listed.

#### Fluorescence Microscopy

Electrode biofilms were visualized using an inverted microscope equipped with UV fluorescence detection. Cells were visualized using the lipid stain FM^TM^ 4- 64FX (Molecular Probes, Life Technologies) at concentrations specified by the manufacturer. RedoxSensor™ Green (RSG) (Molecular Probes, Life Technologies) studies were performed by adding approximately a 10 μM concentration of RSG in addition to lipid stain. Changes in fluorescence activity were monitored over time via microscopy. In brief, biofilm images were taken every five min over a period of six to ten hours depending on the experiment.

Fluorescence intensity between control (open circuit), cathodic (applied reduced voltages), and inhibitor addition conditions were compared using approximately 80 images from each experimental state (including six different time-points taken over a 30 min period). Cellular fluorescence intensities were compared for cells in each image using custom scripts in MATLAB (codes available upon request).

#### ATP, ADP and protein quantification

Electrode biofilms were boiled for 10 min in nanopure water to release cellular ATP and ADP directly from electrode attached cells. For protein recovery from electrode biofilms, samples were treated similarly with the exception that 10mM NaOH was added and samples were heated for at least 1hr at >85°C. Quantification of ATP and ADP was performed using ATP/ADP ratio Assay Kit (Sigma-Aldrich Co.) were performed according to manufacturer’s specification (standard curves for ATP and ADP provided, *Supporting information* Fig. S10). Protein quantification was performed using the NanoOrange^®^ Protein Quantification kit (Molecular Probes, Life Technologies) according to manufacturer’s specifications. Both luminescence and fluorescence were quantified on a BioTek (Winooski, VT) Synergy H4 microplate reader available through the USC Nanobiophysics core facility (dornsife.usc.edu/nanobiophysicscore/).

#### Quantification of *In Vivo* Luciferase Activity

Light emissions from an electrode biofilm were detected across an ITO-coated glass electrode using a photon multiplier tube and associated software (Photon Systems Inc., CA). Uncoupler experiments were performed using either the above systems or a Gene Light 55 GL-100A luminometer (Microtec, Japan). All experiments were conducted in a dark box to minimize light exposure outside of the experiments.

Light emissions for cells grown under various lactate concentrations were quantified in the same reactor setups, however the ITO-coated glass was replaced with a glass microscope slide *(Supporting information* Fig. S3). For these experiments, an LB grown *SO-lux* was diluted into a DM media amended with 7 mM lactate. After ~12 hours of growth cells were rinsed and resuspended at an O.D. 600 of 0.2 in media with varying lactate concentrations (7, 0.7, and >0.07mM lactate). Light production was measured post density normalization and averaged over 20 min time period.

#### Statistical Methods

All experiments were performed in biological replicate (n = 3, unless specified). Statistical analyses were performed using R and/or Excel. In the majority of experiment where quantitative comparisons were made, data were normalized to cell biomass using either protein content (in the case current comparisons for mutants) or total nucleotide pool (in the case of ATP values). In most cases the cell populations for each experiment were constrained by the surface area of the electrode (380 mm^2^) and ensuring surface attachment and planktonic cell removal through repeated washing. Manual cell counts were performed in ImageJ for time zero and time final biofilms for each experiment to ensure that population size remained consistent (20-25% standard deviation from mean value measured for single populations). Twenty images from each biofilm (accounting for 0.1 mm^2^), demonstrated a normal sample distribution around the mean. Results of representative experiments, mean values and standard deviations of biological replicates are reported in figure legend or in text.

## Acknowledgements

Annette Rowe was funded primarily by a Center for Dark Energy Biosphere Investigations (C-DEBI) postdoctoral fellowship, followed by a NASA Astrobiology Institute (NAI) postdoctoral fellowship as part as the NAI Life Underground team. This work is C-DEBI contribution # and NAI contribution #. Part of this work was performed during a JSPS fellowship by Annette Rowe (PE15019) with Dr. Kazuhito Hahimoto at Tokyo University, who we sincerely thank for use of lab equipment. Work in the Nealson lab was funded by the Airforce Office of Scientific Research (GA9550-06-01-0292). Work in the El-Naggar laboratory was supported by Innovation Fund Denmark Electrogas project (partial support for Annette Rowe) and by the Division of Chemical Sciences, Geosciences, and Biosciences, Office of Basic Energy Sciences of the US Department of Energy through grant DE-FG02-13ER16415. Abhiney Jain and Jeffrey Gralnick were supported by the Office of Naval Research (N000141310552). We would like to express our gratitude to Joanna B. Goldberg for supplying us with Lux plasmids from their 2013 paper (26). Thanks to Lina Bird for supplying the bacterial strain BW29427 used for conjugation experiments.

## Supporting Information

Supplementary **Videos S1-S6**. Monolayer MR-1 biofilms treated with a FM-464Fx lipid stain and RSG under various conditions: Aerobic cathode plus cyanide (S1-S3), Aerobic cathode plus Antimycin (S4-S6); **Fig. S1**. Schematic of MR-1 cell inner membrane (A) and outer membrane (B) highlighting cellular components that could interact with cathode-derived electrons; **Fig. S2**. Reduction of oxygen on poised potential cathodes depends on the presence of oxygen and live cells; **Fig. S3.** Light production via MR-1 Lux under aerobic conditions with different concentrations of lactate; **Fig. S4**. Light production under various poised redox potentials in a MR-1 Lux biofilm; **Fig. S5.** Cathodic current during for experiments shown in Fig. 5; **Fig. S6.** Light production on cathode biofilms treated with CCCP with and without lactate present (A) as well as under minimal anodic current conditions (B) and treated with Piercidin A prior to CCCP addition. **Fig. S7.** Cathodic current production normalized to cell biomass in gene deletion mutants of MR-1. **Fig. S8.** Cell decay over time at cathodic current and minimal anodic current producing poised potential; **Fig. S9.** Standard curves for ATP and ADP generated by the luciferase assay.

